# Refinement and Retrospective Shift of the CA1 Place Code during Food-Carrying Decisions

**DOI:** 10.64898/2026.09.14.751532

**Authors:** Zahra Rezaei, Ian Q. Whishaw, Robert J. Sutherland, HaoRan Chang, Majid H. Mohajerani

## Abstract

Food carrying is an adaptive component of foraging that integrates resource evaluation with home-directed navigation, providing a naturalistic model for investigating how the brain organizes goal-directed behaviour. Despite extensive work on hippocampal coding during navigation and homing, how CA1 representations are reorganized when animals spontaneously transport acquired food home remains largely unexplored. We recorded dorsal CA1 population activity using one-photon calcium imaging with a V4 Miniscope while mice performed a self-paced foraging task in which food could be consumed at the acquisition site followed by a homeward return or carried to the home for consumption. Mice preferentially carried larger pellets, indicating that transport behaviour was sensitive to resource value. CA1 spatial coding differed systematically across behavioural outcomes. During carry-homeward runs, spatial representations were sparser, conveyed more spatial information, and showed greater trial-to-trial stability than during eat-inward runs, indicating a more precise and reliable place code during home-directed transport. Place fields also shifted forward during carrying, consistent with a retrospective bias toward recently traversed locations. By linking a self-generated food-handling decision to coordinated changes in the precision, stability, and temporal organization of CA1 activity, these findings extend hippocampal spatial-coding frameworks from trained navigation to ecologically grounded foraging behaviour.

## 2 Introduction

Food carrying is a widespread foraging strategy in which animals transport acquired food from the site of encounter to a safer location for consumption or storage. Whether an item is consumed or carried depends on the value of the resource as well as the costs and risks associated with transport. Across species, larger or more valuable food items are more likely to be transported, whereas increasing distance from protective cover can reduce carrying, reflecting trade-offs among resource value, energetic expenditure, and predation risk (Lima et al., 1985; Lima and Valone, 1986; Valone and Lima, 1987). In rodents, food size and anticipated eating time strongly influence whether food is consumed where it is found or transported to a refuge, and larger items can elicit shorter carrying latencies, faster transport and longer return to search (Whishaw, 1990; Whishaw et al., 1990). Carrying is further reduced as travel distance or effort increases (Whishaw and Dringenberg, 1991). Once carrying is initiated, the return to the home base is typically rapid and direct relative to the more variable outward journey (Whishaw et al., 2001; Wallace et al., 2002). The hippocampal formation appears to contribute to this organization. Hippocampal damage does not necessarily abolish food carrying, but disrupts its adjustment to environmental change and increasing travel distance (Whishaw, 1993), while disruption of hippocampal pathways impairs the directness and accuracy of homeward navigation (Whishaw et al., 2001; Wallace et al., 2002). Moreover, hippocampal populations encode elapsed time and travelled distance during ongoing experience (MacDonald et al., 2011; Kraus et al., 2013), providing a potential substrate for integrating spatial and temporal variables that strongly influence food-handling decisions. Together, these observations suggest that food carrying recruits processes that link resource evaluation with the spatial and temporal organization of a directed return to home.

There are many reasons to propose that hippocampal cellular representations could contribute to foraging behavior. Their collective organization associated with cognitive-map framework (Tolman, 1948; O’keefe and Nadel, 1979). Spatial firing varies with internal and behavioural state (Kennedy and Shapiro, 2009), and place fields can become concentrated around goal locations (Hollup et al., 2001) and reward sites (Hok et al., 2007). CA1 activity can also distinguish intended destinations (Ainge et al., 2007) and routes through otherwise overlapping space (Grieves et al., 2016), indicating that motivational and contextual information is incorporated into hippocampal representations during goal-directed navigation (Redish, 2016; Wikenheiser and Schoenbaum, 2016). Goal-directed signals can directly reshape spatial firing during navigation (Aoki et al., 2019; Ormond and O’Keefe, 2022), and hippocampal population representations can vary according to the relationship between the origin and destination of a journey (Gothard et al., 1996). CA1 recordings during a path-integration-based task revealed distinct representations during search and homing, with task-anchored firing fields carrying information predictive of homing direction (Najafian Jazi et al., 2023). Furthermore, hippocampal ensembles contribute to goal-directed behaviour through prospective and retrospective coding. Prospective representations can encode future trajectories and remembered goals (Pfeiffer and Foster, 2013; Sarel et al., 2017), whereas retrospective representations can preserve information about recently visited locations and preceding behavioural states (Ferbinteanu and Shapiro, 2003; MacDonald et al., 2011). Place fields themselves can shift along or opposite the direction of travel with experience (Mehta et al., 1997; Lee et al., 2004, 2006; Battaglia et al., 2004; Xu et al., 2019), consistent with dynamic weighting of recently experienced and upcoming locations according to behavioural demands (Shin et al., 2019). The stability and precision of hippocampal representations are similarly sensitive to behavioural significance. Attention to spatial context increases place-field stability (Kentros et al., 2004), while goal-and reward-related learning can reorganize and stabilize behaviourally relevant place-cell ensembles (Hok et al., 2007; Dupret et al., 2010; Danielson et al., 2016). These properties make the hippocampal place code well suited to differentiate behavioural states that occur within the same physical environment but differ in their goals, motivational significance, and relationship to recent experience. Although hippocampal activity has been characterized during goal-directed navigation and homing, how these coding properties are reorganized during the spontaneous transport of an acquired reward remains poorly understood.

Here we examined whether the complexity of food carrying including resource value, food size, distance to home, hunger state, food-handling time, and risk exposure (Bindra, 1947, 1948b,a; Ewer, 1971; Lima et al., 1985; Lima and Valone, 1986; Whishaw, 1990; Whishaw and Dringenberg, 1991) influence hippocampal cellular activity. Given these spatial, temporal, and motivational demands, we hypothesized that carrying food home would engage a distinct hippocampal coding state compared with returning after consuming food at the acquisition site. Specifically, we predicted that carrying would be associated with more precise and stable CA1 spatial representations and with a systematic reorganization of place-field position relative to outward travel. To test these predictions, we recorded dorsal CA1 population activity using one-photon calcium imaging while mice performed a self-paced foraging task in which acquired food could either be consumed at the food site or carried to a home compartment. By characterizing CA1 population dynamics during spontaneous food carrying vs returning without food, our study provides a neural account of an ecologically meaningful foraging behaviour that integrates resource evaluation with home-directed navigation, extending hippocampal research beyond trained tasks to the organization of self-generated, adaptive behaviour.

## 3 Results

### 3.1 Food size biases spontaneous carrying decisions

We first established whether mice expressed spontaneous food carrying in our foraging task and whether the decision to carry was sensitive to food size. Mice retrieved food pellets from an alley and could either consume them at the food site or carry them back to the home compartment (Figure 1c) (see Methods 6.5). The proportion of successful trials (i.e. EAT or CARRY) remained largely stable across days (Figure 1d). A mild habituation phase was observed, with performance stabilizing around day five. To confirm that performance chart consists of two phases, regression analysis was performed on each part separately. Piecewise mixed-effects regression revealed distinct dynamics between early and late phases. During the early phase (Days 1–5), behaviour changed significantly, as indicated by a nonzero slope (*β* = 0.098*, p* = 1.86 × 10*^−^*^10^). In contrast, during the late phase (Days 6–32), the slope did not significantly differ from zero (*β* = −0.001*, p* = 0.453), indicating stable performance after initial habituation. These findings suggest that task adaptation occurs primarily in the first few days, followed by a plateau phase. This consistency suggests that the task relies on innate foraging strategies rather than requiring extended learning or training.

**Figure 1:**
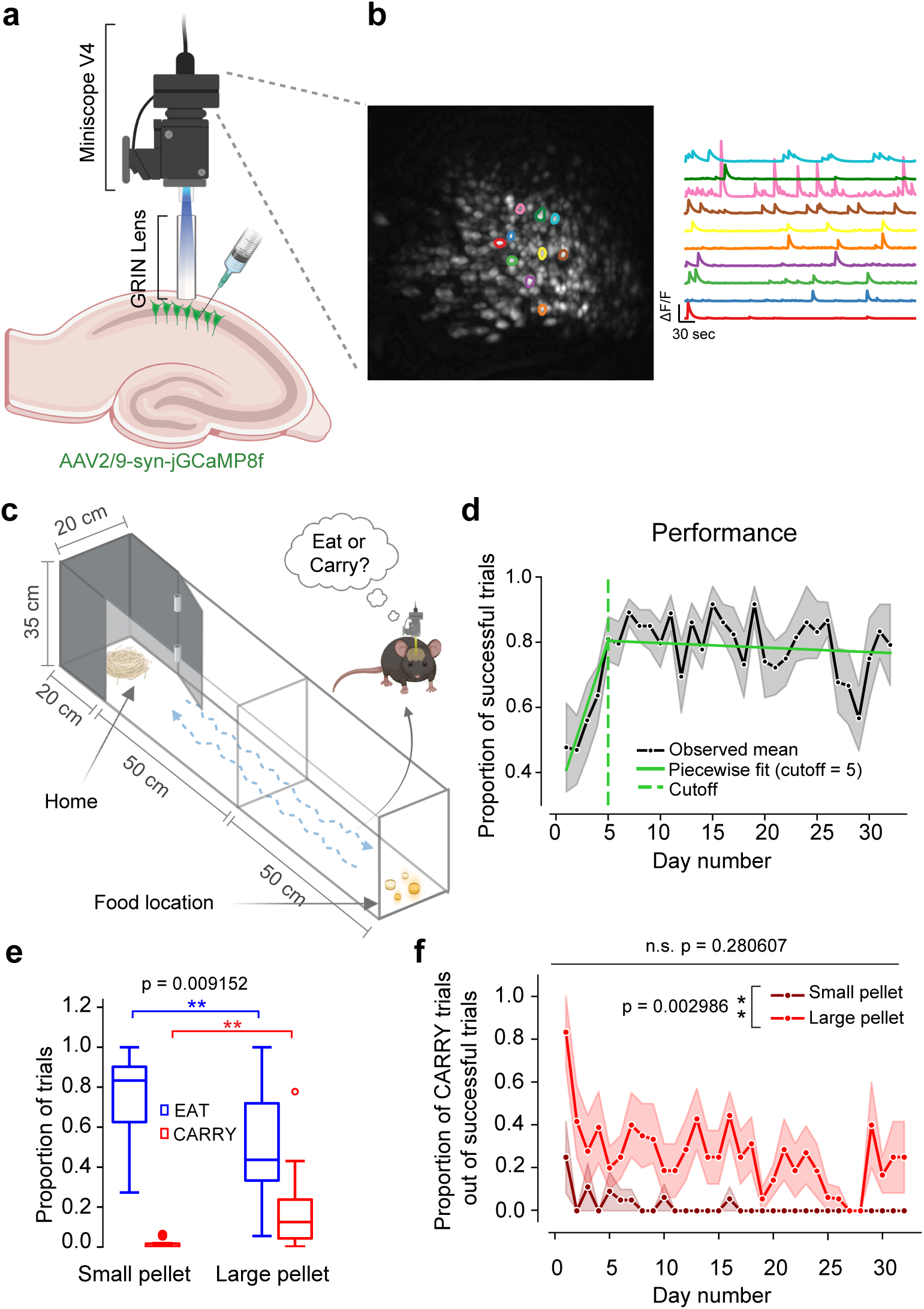
Experimental design, calcium imaging, and behavioural performance in the foraging task. **a** Schematic of viral injection and GRIN lens implantation for *in vivo* calcium imaging with a Miniscope (see Methods 6.6). **b** Example field of view showing putative dorsal CA1 pyramidal cells with corresponding fluorescence time traces. **c** Behavioural arena used for the foraging task. Experiments were conducted with alleys of two different lengths: 50 and 100 cm. Food pellets were prepared in four sizes (see Methods 6.2, 6.4, and 6.5). **d** The proportion of successful trials over experimental days. A piecewise linear mixed-effects model was used to assess changes across early (Days 1–5) and late (Days 6–32) phases. The model revealed a significant slope in the early phase (*p* = 1.8 × 10*^−^*^10^), but no significant change in the late phase (*p* = 0.45). **e** The proportions of EAT vs CARRY trials out of all successful trials for small and large food pellets. Pellet size significantly affected behavioural strategy: eating decreased while carrying increased for large pellets (paired-sample Wilcoxon signed rank test was performed on the CARRY condition only, since there is only one degree of freedom, i.e. %CARRY = 100% − %EAT; *p* = 0.009152). **f** The proportion of CARRY trials among successful trials across days. A linear mixed model was fitted to investigate the effects of pellet size and exposure day on the proportion of CARRY trials (R lmer formula: proportion ∼ Day ∗ PelletSize +(Day ∗ PelletSize|MouseId)). Type III ANOVA with Satterthwaite’s method showed a main effect for pellet size (*F* (1, 10.3582) = 14.8613; *p* = 2.986 × 10*^−^*^3^), but not for days (*F* (1, 9.9707) = 1.3015; *p* = 0.280607). A significant interaction was observed (*F* (1, 14.1776) = 6.3298; *p* = 0.024516).

Pellet size strongly influenced foraging strategy. The probability of eating at the food site decreased for large pellets, while the probability of carrying them back to the refuge increased (Figure 1e). This trade-off between local consumption and transport was statistically significant (Wilcoxon signed rank test, *p* = 0.009152), highlighting the role of energetic or handling constraints in shaping decision-making. Importantly, pellet size exerted a main effect on the proportion of carry responses within successful trials, whereas day number had no significant effect (Figure 1f). A linear mixed model was fitted to investigate the effects of pellet size and exposure day on the proportion of CARRY trials (R lmer formula: proportion ∼ Day∗PelletSize+(Day∗PelletSize|MouseId)). Type III ANOVA with Satterthwaite’s method showed a main effect for pellet size (*F* (1, 10.3582) = 14.8613; *p* = 2.986 × 10*^−^*^3^), but not for days (*F* (1, 9.9707) = 1.3015; *p* = 0.280607). A significant interaction was observed (*F* (1, 14.1776) = 6.3298; *p* = 0.024516). Thus, the observed behavioural variation was driven by food-related factors rather than task exposure or experience, therefore underscoring an instinctual form of decision-making rather than an acquired behaviour.

### 3.2 Food carrying sharpens and stabilizes CA1 place coding

We next tested whether carrying food home was associated with a distinct and more precise CA1 spatial representation, as predicted from the structured and goal-directed nature of the homeward journey. Because these analyses treated the foraging alley as a one-dimensional environment, we first verified that animal movement was dominated by the longitudinal axis of the apparatus. Positional variance along the longitudinal axis was approximately 92-fold greater than variance along the lateral axis, supporting linearization of position for subsequent analyses (Figure S1b).

Using this linearized representation, we identified neurons with persistent spatial tuning across outgoing runs, inward runs following food consumption (EAT), and inward runs while carrying food (CARRY) (Figure 2a; Figure 3a). Although many neurons maintained spatially localized responses across conditions, their tuning differed markedly during carrying. In particular, CARRY trials showed a pronounced reduction in out-of-field activity compared with the other conditions (Figure 2a; Figure 3a). At the population level, this was accompanied by a steeper decline in population-vector correlation with spatial displacement (Figure 2b), indicating greater spatial differentiation of CA1 population activity during food carrying. To quantify this refinement at the single-cell level, we measured the lifetime sparsity of neuronal tuning curves across conditions. Tuning was sparsest during CARRY trials, supporting greater spatial selectivity during food carrying (Figure 2c). EAT trials also showed increased sparsity relative to outgoing trials, although the effect was smaller than during CARRY trials. Quantification of spatial information revealed a similar trend, wherein place cells conveyed significantly higher spatial information on CARRY trials, while a modest increase in spatial information was noted on EAT trials compared to outgoing trials (Figure 2d). In consideration of the low number of trials inherent to the nature of the behavioural task, we employed an alternative method for computing spatial information that is less susceptible to bias from sample size (see Methods 6.9). Interestingly, a decoder trained on the ingoing trials exhibited higher accuracy in positional inference when decoding locations during outgoing trials than the other way around, further underscoring an overall more refined spatial population code on ingoing trials (Figure 3d). Finally, trial-to-trial stability, quantified as the mean Pearson correlation between spatial tuning curves across trial pairs, was significantly higher during CARRY than EAT trials (Figure 2e).

**Figure 2:**
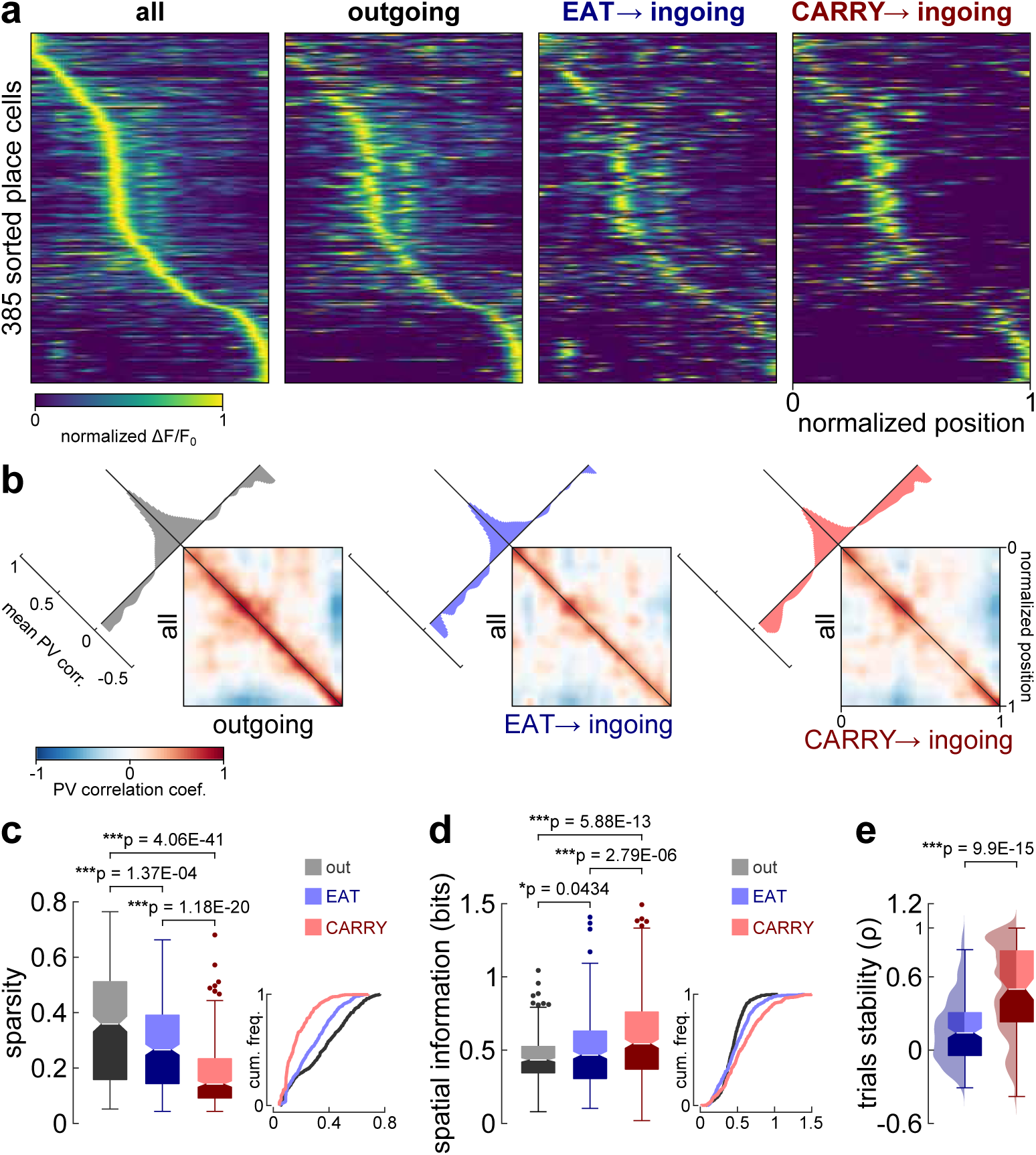
Heightened precision and stability in place coding during the carrying of food. **a** Normalized Δ*F/F*_0_ as a function of position for all identified place cells (see Methods 6.8) averaged over: 1. all trials and directions, 2. outgoing direction (from home to goal), 3. EAT trials on ingoing direction, and 4. CARRY trials on ingoing direction. Place cells are sorted by location of peak response across all trials. **b** Correlation matrices of population vectors (PV) between all trials and individual trial conditions. Inset shows average correlation coefficient as a function of shift from the central diagonal. **c-d** Sparsity and spatial information of place cells on: 1. outgoing trials, 2. ingoing EAT trials, and 3. ingoing CARRY trials. Kruskal-Wallis tests showed a significant difference across the medians of trial conditions in both sparsity (*p* = 1.9249 × 10*^−^*^42^) and spatial information (*p* = 6.7095 × 10*^−^*^13^). Post-hoc multiple comparisons on mean ranks were subsequently performed, and p-values were adjusted by the Bonferroni method. Insets depict data as cumulative frequency. **e** Mean stability (correlation coefficient) of spatial responses across trials for the EAT and CARRY conditions. A two-sample Wilcoxon rank sum test was performed to compare differences in median stabilities. All boxplots show median (line), first and third quartiles (box), minimum and maximum values (whiskers), and outliers (dots).

**Figure 3:**
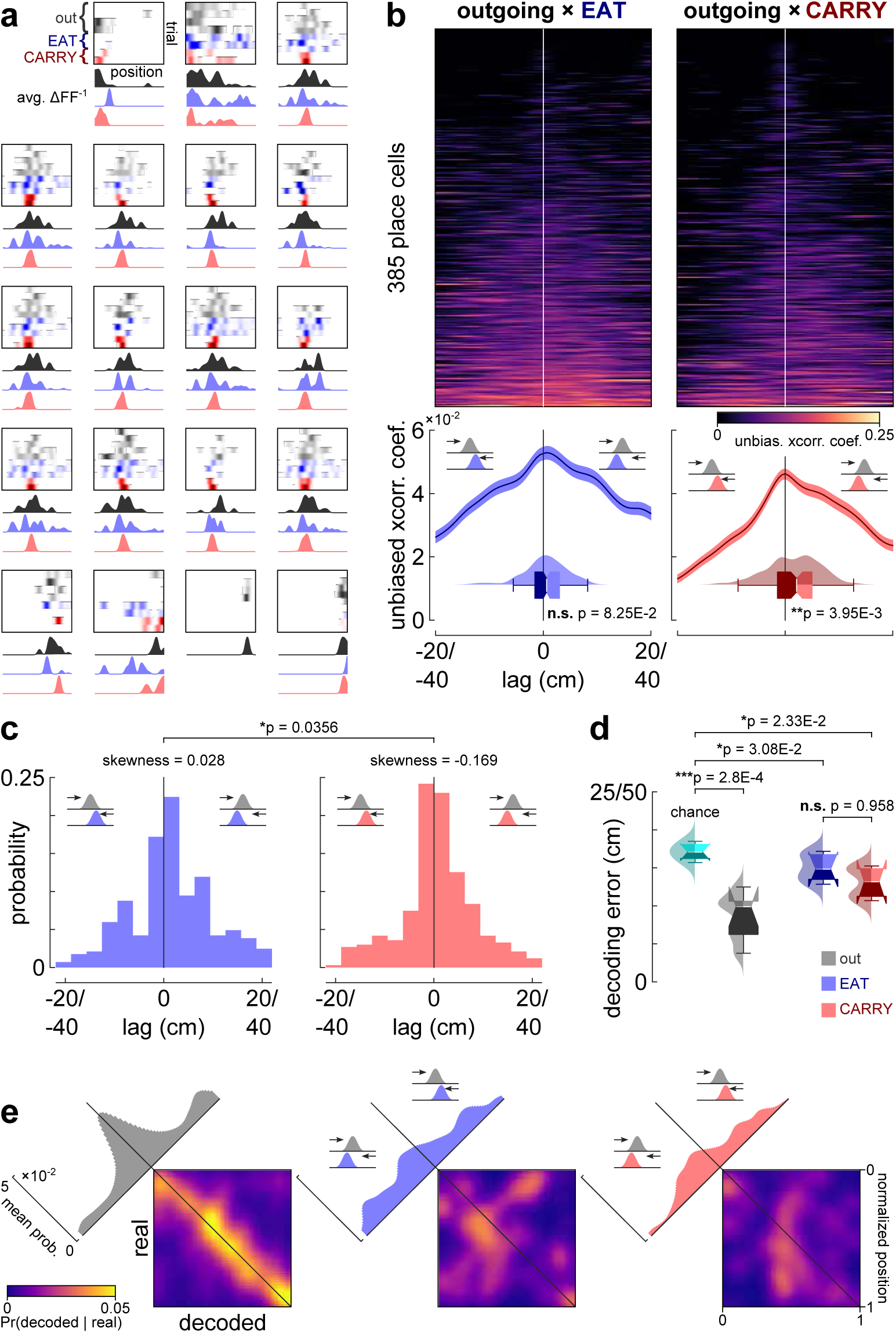
Retrospective shifting of place fields during the carrying of food. **a** Representative examples of place cells across trial conditions. Each box depicts an individual place cell’s response as a function of location on outgoing (black), EAT (blue) and CARRY (red) trials. Averaged responses across trial conditions are shown below. **b** Unbiased cross-correlation coefficients of place cells’ spatial tuning curves between outgoing and EAT/CARRY trials. Neurons sorted by average correlation coefficient. Mean cross-correlation ± SEM are shown below. Boxplots and densities represent each neuron’s average lag distance weighted by the cross-correlation coefficients. Wilcoxon signed rank tests were performed to compare the median lag distances against 0 lag. **c** Distribution of peak cross-correlation lag distances for place cells’ tuning curves between outgoing and EAT/CARRY trials (see b). The distribution for CARRY trials shows a stronger negative skew than EAT trials (bootstrap test for equal skewness), suggesting a stronger tendency in forward shifting on CARRY trials. **d** Average position decoding error (measured in absolute distance) for different trial conditions (outgoing runs, EAT ingoing and CARRY ingoing runs). Only recording sessions containing more than 30 place cells were analysed (*n* = 7 recording sessions). For outgoing trials, the decoder was trained on all ingoing trials, while for EAT and CARRY ingoing trials, the decoder was trained on all outgoing trials. Decoding error was significantly lower than the theoretical chance level for all conditions, while accuracy was comparable between EAT and CARRY (paired-sample t-tests). **e** Confusion matrices representing the probability of decoded positions conditioned on the real positions, averaged across all 7 sessions. Inset shows average probability as a function of shift from the central diagonal.

Because differences in movement paths could potentially influence apparent spatial tuning, we performed a complementary trajectory-control analysis (Figure S1c-e and Table S1). We compared RMS lateral deviation and tortuosity across outgoing, EAT, and CARRY trajectories separately for the short and long alleys (see Supplementary). Although some comparisons showed large matchedpairs effect sizes, none of the comparisons remained significant after Holm correction. Thus, the condition-dependent differences in CA1 coding are unlikely to be explained solely by gross differences in trajectory geometry. Together, these results support our prediction that food carrying is accompanied by a more precise and stable CA1 spatial representation.

### 3.3 Retrospective shift in place fields during food carrying

We next tested our prediction that carrying would alter the spatial organization of place fields relative to outward travel. All spatial analyses were performed in the global reference frame of the maze, such that position corresponded to absolute linearized location along the track rather than distance from the starting point of each run. Within this reference frame, comparison of spatial tuning profiles revealed an overall shift of place fields toward home on CARRY trials relative to outgoing trials, corresponding to a forward displacement along the inward direction of travel (Figure 3a-b). This forward displacement reflects a retrospective coding bias, as place-cell activity occurred later along the trajectory, after the animal had passed the location represented by the corresponding outgoing field. Cross-correlation analysis confirmed a significant retrospective shift between outgoing and CARRY tuning curves, whereas no significant shift was detected between outgoing and EAT trials (Figure 3b). The distribution of peak cross-correlation lags was also more negatively skewed for CARRY than EAT trials (Figure 3c), further supporting a carrying-specific retrospective bias. At the population level, a decoder trained on outgoing activity similarly assigned CARRY activity to positions displaced toward recently traversed locations, whereas no comparable bias was evident for EAT trials (Figure 3d-e). Together, these results indicate that food carrying reorganizes the spatial alignment of CA1 place fields and biases the population code toward recently experienced locations during the homeward journey.

## 4 Discussion

We employed a self-paced foraging paradigm in which mice decided whether to consume or carry food pellets to a home base, thereby engaging a naturalistic decision-making scenario. We found that larger pellets were preferentially carried rather than eaten in place, and that hippocampal spatial representations during carrying exhibited greater precision and stability than during returns without food. Furthermore, place fields shifted forward along the return path during carrying, consistent with a retrospective coding bias. These findings support our prediction that food carrying engages a distinct hippocampal coding state and illuminate how CA1 spatial representations adapt to motivational context, goal-directed behaviour, and the intrinsic decision of whether to consume or transport an acquired resource.

### 4.1 Food carrying as naturalistic decision-making

Food carrying provides a particularly useful setting for studying naturalistic decision-making because the choice to consume or transport an acquired resource emerges within an ongoing foraging episode rather than through extensive training. In our paradigm, mice were not taught an arbitrary response rule after encountering food. Instead, they spontaneously chose whether to eat at the food site or carry the resource home. Carrying is itself an adaptive strategy for relocating food to locations where it can be consumed or stored under more favourable conditions (Andersson and Krebs, 1978; Smulders et al., 2010). Because carrying behaviour is shaped by resource value, handling demands, travel cost, and risk, it preserves the ecological structure of the decision while allowing its neural correlates to be examined under controlled conditions (Mobbs et al., 2018).

Consistent with classic food-carrying studies, mice in our task preferentially transported larger pellets, while exposure day had no comparable main effect after the initial habituation period. Previous work has similarly shown that carrying increases with food size or anticipated handling time and is adjusted according to distance, effort, food availability, and risk (Lima and Valone, 1986; Lima et al., 1985; Whishaw, 1990; Whishaw and Dringenberg, 1991). The stability of this behaviour across days, together with its continued sensitivity to pellet size, supports the interpretation that carrying reflects an intrinsic food-handling strategy rather than gradual acquisition of a task-specific choice rule. This provides the behavioural context for interpreting the carry-related reorganization of hippocampal coding described below.

### 4.2 Hippocampal spatial coding: precision, stability, and goal context

The hippocampus represents more than an animal’s physical location. Its spatial activity is shaped by the motivational and contextual significance of ongoing experience (Ainge et al., 2007; Hok et al., 2007; Hollup et al., 2001; Kennedy and Shapiro, 2009; O’Keefe and Krupic, 2021; Redish, 2016; Wikenheiser and Schoenbaum, 2016). Hippocampal spatial selectivity is sensitive to the structure of behaviour, becoming weaker during passive movement and more precise when navigation is organized around meaningful goals (Aghajan et al., 2015). Goal-related information can be incorporated directly into the spatial map, with movement toward a goal enhancing in-field firing and spatial coherence (Aoki et al., 2019), while route-dependent coding indicates that hippocampal representations reflect the unfolding behavioural sequence rather than destination alone (Grieves et al., 2016). Within this framework, the greater precision and stability observed during carrying suggest that food transport engages a coherent and behaviourally meaningful navigational state. Although no carry and carry returns involved inward movement toward home, carrying required the animal to transport a valuable resource toward a consistent and salient destination, potentially strengthening the organization of the hippocampal representation (Aoki et al., 2019; Grieves et al., 2016; Ormond and O’Keefe, 2022).

Place-field stability is an important feature of reliable spatial representation and varies with the behavioral relevance of the environment and task demands (Kentros et al., 2004; Zhang and Manahan-Vaughan, 2015). Stable hippocampal representations are associated with successful spatial learning, and reward-related learning can selectively reorganize and stabilize behaviourally relevant place-cell ensembles (Danielson et al., 2016; Dupret et al., 2010). Reward expectation may provide one explanation for the enhanced stability during carrying. Removal of an expected reward degrades CA1 spatial maps, whereas stronger reward expectation is associated with more reliable representations and reduced representational drift (Kaufman et al., 2020; Krishnan et al., 2022; Krishnan and Sheffield, 2023). Thus, carrying an acquired resource toward home likely increases the motivational significance of the return journey relative to inward travel after the resource has already been consumed.

A complementary possibility is that carrying places greater demands on dead reckoning, or path integration, during the homeward journey. Dead reckoning uses accumulated self-motion information to update position relative to a reference location and can support direct return to home when external cues are limited and is supported by the hippocampus (Mittelstaedt and Mittelstaedt, 1980; Whishaw et al., 2001). Food-carrying and exploratory homing studies have implicated the hippocampal formation in this process, showing that hippocampal disruption alters the directness and accuracy of homeward navigation (Whishaw et al., 2001; Wallace et al., 2002). More recently, CA1 representations were shown to reorganize between search and homing phases during path-integration-based behaviour, with hippocampal firing fields carrying information predictive of homing direction (Najafian Jazi et al., 2023). Carrying toward a fixed home location may therefore engage a more consistent integration of self-motion and goal-related information, potentially contributing to the enhanced stability of the CA1 representation. Carrying food toward a fixed home location may engage a consistent integration of self-motion, reward value, goal approach, and organized action sequences, thereby contributing to the sharpening and stabilization of CA1 spatial representations.

### 4.3 Forward shift of place fields: retrospective encoding

Shifts in hippocampal place fields are a well-established feature of spatial coding, and their functional significance depends on the behavioural context in which they occur. For example, backward shifts relative to the direction of travel are associated with prospective or anticipatory coding of upcoming locations via experience-dependent plasticity and theta-phase dynamics (Mehta et al., 1997; Battaglia et al., 2004). Forward translocations, although less commonly reported, have also been observed during goal-directed learning, with fields progressively shifting toward prospective reward locations (Lee et al., 2006; Xu et al., 2019). In our task, place fields shifted forward toward home during inward carry trials, whereas no comparable shift was observed during returns without food. During these homeward carrying runs, the forward displacement biased neuronal activity toward locations the animal had recently traversed, consistent with retrospective coding. Such retrospective or history-dependent hippocampal representations have been reported in other task contexts, in which neuronal activity retains information about recently traversed routes, journey origin, or preceding behavioural states (Ferbinteanu and Shapiro, 2003; Catanese et al., 2014).

The retrospective bias may reflect the particular mnemonic structure of food carrying. Retrospective and prospective hippocampal representations can coexist within the same environment, with their relative expression shaped by task demands and behavioural state (Ferbinteanu and Shapiro, 2003; Catanese et al., 2014). Replay dynamics similarly distinguish past from future trajectories, with reverse replay linked to recently completed paths and forward replay to prospective trajectories (Shin et al., 2019). During carrying, the recently visited food location and the route leading to it may remain relevant because acquisition and transport form a continuous foraging episode that is not completed until the resource reaches home. The homeward journey may therefore maintain or reactivate information about the immediately preceding path more strongly than inward travel after food has already been consumed.

Several complementary mechanisms could contribute to this carry-related shift. One possibility involves reward-associated reverse replay and subsequent modification of hippocampal sequential representations. Recently traversed trajectories can be replayed in reverse order following reward or spatial experience (Lee and Wilson, 2002; Foster and Wilson, 2006), and reverse activation has been proposed to alter previously established sequential associations in a manner that can produce forward translocation of place fields (Lee et al., 2006; Ponzi, 2009). A complementary mechanism may involve persistence of recent positional information within entorhinal/postrhinal–hippocampal circuits, allowing representations of preceding locations to overlap with incoming spatial information (Lee et al., 2006). During food carrying, either process could strengthen the influence of the recently completed food-directed trajectory on the evolving homeward representation, providing a potential basis for the retrospective bias observed here.

### 4.4 Limitations and future directions

Several limitations should be considered when interpreting these findings. The naturalistic, self-paced design yielded relatively few trials per recording session, limiting statistical power and the extent to which behavioural variables could be matched across conditions. Although complementary trajectory analyses suggested that gross differences in path geometry were unlikely to account for the observed neural effects, subtler behavioural contributions cannot be excluded. Place cells were also identified using trials pooled across behavioural conditions and movement directions to preserve statistical power, and the present study therefore did not address possible direction-dependent remapping. Finally, one-photon calcium imaging did not provide the temporal resolution or electrophysiological signals required to directly examine theta-related activity, sharp-wave ripples, or fast replay dynamics, and path integration was not experimentally manipulated. Accordingly, the dead-reckoning and replay-related mechanisms discussed above remain plausible interpretations rather than processes directly demonstrated here.

Future studies combining cellular-resolution imaging with electrophysiological recordings could determine whether theta dynamics or replay accompany the carry-related refinement and retrospective shift of the CA1 place code. Manipulating visual and self-motion cues during the homeward journey would provide a more direct test of the contribution of dead reckoning. Extending recordings to connected regions, including entorhinal, retrosplenial, prefrontal, and striatal circuits, could further clarify how hippocampal spatial representations interact with navigation, valuation, and action selection during naturalistic food carrying.

## 5 Conclusion

In summary, food carrying was associated with a distinct CA1 spatial code during naturalistic foraging. When mice transported food toward home, spatial representations became more precise and stable, and place fields shifted forward along the homeward trajectory, producing a retrospective bias toward recently traversed locations. These effects were not readily explained by gross differences in trajectory geometry, supporting the interpretation that the hippocampal representations are reorganized according to the behavioural and motivational significance of an ongoing foraging action rather than physical position alone. By linking the intrinsic decision to consume or transport an acquired resource with coordinated changes in the precision, reliability, and temporal organization of CA1 activity, our findings extend the hippocampal cognitive-map framework to the neural organization of naturalistic goal-directed behaviour.

## 6 Methods

### 6.1 Animals

Ten C57 mice (six males and four females) were housed individually in cages from two months of age under a 12-h light–dark cycle. At five months of age, the animals were handled daily for one week and then subjected to food restriction. Once they reached 85% of their baseline body weight, they were introduced to the task environment for approximately one hour per day for a week. The body weight maintained at 85% of the baseline weight throughout the experiments which was the lowest allowed body weight by the protocol. After a week, surgery was performed. Food restriction was lifted one day prior to surgery and remained suspended for two weeks postoperatively. All animal procedures were approved by the University of Lethbridge Animal Care Committee and were conducted in accordance with the guidelines of the Canadian Council on Animal Care.

### 6.2 Food

Four sizes of food pellets were used: 14 mg (0.125”), 45 mg (0.156”), 190 mg (0.175”), and 300 mg (0.281”). All pellets were Dustless Precision Pellets (Bio-Serv) and were unflavoured. During experimental sessions, only pellets were provided. Whereas, in the home cage, mice had access to both standard facility chow and pellets, supplied at a 70:30 ratio.

### 6.3 Surgical procedure

The surgery procedure consisted of four main stages.

#### 6.3.1 Craniotomy and tissue aspiration

Mice underwent cortical aspiration to expose the CA1 hippocampus for GRIN lens implantation (Resendez et al., 2016). A craniotomy (2 mm posterior, 2 mm lateral to bregma; 1 mm radius) was performed under isoflurane anaesthesia (induction: 4–5% with 4 L/min O_2_; maintenance: 1.5–2% with 1 L/min O_2_) with the head secured in a stereotaxic frame and body temperature maintained at 37 °C. The skull was drilled under sterile saline irrigation, and gel foam was applied to the exposed tissue. Aspiration was performed with a bent 27G needle connected to a vacuum pump while continuously irrigated with saline. Bleeding was controlled with gel foam. Aspiration was stopped after passing the corpus callosum and reaching the alveus, leaving a thin tissue layer above CA1. The site was then filled with sterile saline.

#### 6.3.2 AAV injection

Following cortical aspiration, Adeno-Associated Virus (AAV) injected to introduce the Ca^2+^ indicator into the neural population (Resendez et al., 2016). For this purpose, we used the GCaMP8f viral vector, which has a nearly tenfold faster fluorescence rise time than previous GCaMPs and can track individual spike rates up to 50 Hz (Zhang et al., 2023). A Nanoject injector was loaded with 400 nl of AAV2/9-Syn-jGCaMP8f (titer 1.6 × 10^13^) and positioned above the aspirated site (1.3 mm below the brain surface). Virus was infused at 50 nl/min at four sites, each 250 µm medial–lateral and anterior–posterior to the lens centre. After infusion, the needle remained in place for 5 min to allow diffusion, then was withdrawn slowly at 50 nl/min.

#### 6.3.3 GRIN lens implantation

A 1-mm diameter GRIN lens was implanted into the craniotomy above CA1. The lens was lowered until brain tissue and vasculature were visible, then secured to the skull with Wet-Bond, followed by Metabond. Dental acrylic cement was applied in two layers, the second mixed with carbon powder to reduce light reflection. A Parafilm cap was placed over the lens and sealed with Kwik-Sil for protection.

#### 6.3.4 Baseplate and Miniscope attachment

Three weeks after lens implantation, once GCaMP expression was sufficient, mice underwent baseplate attachment. Under isoflurane anesthesia, Kwik-Sil and Parafilm were removed, and the GRIN lens was cleaned with 70% ethanol. The Miniscope was positioned over the lens using the stereotaxic arm, and the focal plane was adjusted until vasculature and cell bodies were clearly visible. To compensate for shrinkage of dental cement, the Miniscope was raised 50 µm before securing the baseplate with carbon-mixed dental cement. After curing, the Miniscope body was detached, and a protective cap was placed over the baseplate.

### 6.4 Apparatus

Behaviour was tested in a rectangular box (120 × 20 × 35 cm, L×W×H) consisting of a black acrylic refuge (20 × 20 cm) and a transparent Plexiglas alley (up to 100 cm). A sliding end wall allowed the alley length to be set at 50 or 100 cm. Food pellets were placed at the end of the alley.

A gate, controlled by a 90° servo motor, separated the refuge from the alley and opened at trial onset. Nesting material for each animal was placed in the refuge during experiments. Behaviour was recorded using top-and front-view Raspberry Pi cameras connected to an Arduino controller. The UCLA Miniscope V4 (Cai et al., 2016) was used for calcium imaging, interfacing with a DAQ Box and acquisition computer. Synchronization between imaging and behavioural recordings was achieved by sending a signal from the DAQ Box to the Arduino, which triggered both cameras. Recording sessions were initiated and terminated through the Miniscope DAQ software.

### 6.5 Behaviour: foraging task

The task was performed under multiple environmental and reward conditions. Specifically, mice foraged in two alley lengths (50 cm and 100 cm), with food pellets differing in size (four different sizes). Each mouse completed approximately 32 daily sessions in total, with 16 sessions for each distance condition. Because distance did not affect any of the behavioural or brain imaging results, data across the two distance conditions were combined for subsequent analyses. Each daily session consisted of four trials, in which food pellets of four different sizes (see Methods 6.2) were presented in a randomized order. Since behavioural results showed similar patterns between the two smaller and the two larger sizes, data were collapsed into two categories, small and large pellets, for subsequent analyses. Each trial began with the opening of a gate connecting the home compartment to the foraging alley and ended when the animal returned home, at which point the gate was closed. Upon encountering the food pellet, mice exhibited one of three primary foraging behaviours: Eat, Carry, or Nose-poke. In some trials, animals remained in the home compartment (No-leave). Trials resulting in Eat or Carry outcomes were considered successful, while those ending in Nose-poke or No-leave were classified as failed. Each successful trial comprised an outgoing run from the home compartment to the food location followed by an inward return to home. For neural analyses, these movement epochs were treated separately. Outgoing runs were pooled across subsequent behavioural outcomes, whereas inward runs were classified as EAT or CARRY according to whether the animal consumed the food at the acquisition site or transported it home. All spatial analyses were performed in a fixed apparatus-centered coordinate system, with position defined as absolute linearized location along the alley rather than distance from the starting point of an individual run. Additional analyses assessing spatial linearization and trajectory geometry are described in the Supplementary Information (Figure S1 and Table S1).

### 6.6 One-photon microscopy

Calcium imaging was performed using a UCLA Miniscope V4 (Cai et al., 2016; Shuman et al., 2020) coupled with a gradient refractive index (GRIN) lens to record neural activity from pyramidal neurons in the CA1 region of the hippocampus. A GRIN lens (1 mm diameter) was implanted above dorsal CA1 following standard stereotaxic procedures, providing optical access to the underlying neuronal population. After recovery, the Miniscope V4, equipped with a CMOS sensor and integrated excitation LED (peak emission 470 nm), was mounted onto a baseplate affixed to the skull, enabling head-mounted *in vivo* imaging during behaviour. The system provided a field of view of approximately 700 × 450 µm at single-cell resolution and data were acquired at 30 Hz, allowing robust recording of calcium transients from CA1 pyramidal neurons in freely moving mice.

### 6.7 Imaging pre-processing

Calcium imaging data were preprocessed using Minian (Dong et al., 2022). Motion correction, background subtraction, and source extraction were performed using a constrained non-negative matrix factorization (CNMF) framework that jointly estimates spatial footprints and temporal activity of individual neurons. The temporal component was deconvolved using an autoregressive model to infer underlying spiking activity from calcium fluorescence. The resulting deconvolved activity traces (S) were used for all subsequent analyses.

### 6.8 Place cells detection

Two criteria were used for the detection of place cells. First, a place cells must exhibit stable tuning curves over space across trials. For *T* trials, stability for a neuron is measured, here, as the average Pearson correlation coefficient between the response over spatial locations across all trial pairs. That is,

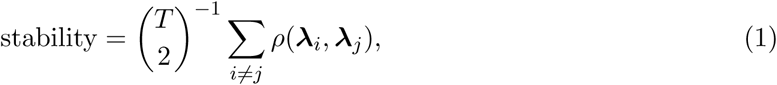

where *ρ*(·) is the Pearson correlation coefficient, ***λ****_i_* is the neuron’s tuning curve on trial *i* and 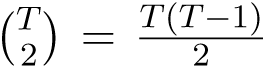 is the number of unique combinations of trial pairs for *T* trials. We compare this stability against a shuffled sample by circularly shifting the tuning curves by a random integer for each trial and computing the stability. This permutation is performed 500 times and neurons whose stability is higher than the 95^th^ percentile of the shuffled samples are kept as candidate place cells. The second criterion requires neurons’ spatial information (see Methods 6.9) to exceed the 95^th^ percentile of shuffled samples. Here, the shuffling procedure involves circularly shifting the Δ*F/F*_0_ time vectors of neurons and recomputing the spatial information. For place-cell detection, stability and spatial information were calculated across trials pooled over behavioural conditions and movement directions (outgoing, EAT-inward, and CARRY-inward). This approach was used to preserve statistical power given the limited number of trials per condition. Hippocampal place cells can exhibit direction-dependent firing on constrained trajectories (Markus et al., 1995; Battaglia et al., 2004), but the present analysis was not intended to identify direction-specific place-cell populations.

### 6.9 Estimation of spatial information

The present work does not follow the standard approach for computing spatial information originally proposed by Skaggs et al. (1992), but instead implements an alternative algorithm. Spatial information is defined as the mutual information between an agent’s spatial location and a neuron’s firing rate. In the original article, this measure was estimated by approximating a neuron’s firing rate as a function of location (typically done through binning), subject to the condition that the firing rate is greater than 0 at all locations. However, due to the sparse nature of both calcium imaging and the design of our task, measuring spatial information in our data using this standard approach leads to severe biases. On the one hand, calcium transients are sparse events requiring a considerable number of trials to obtain a reasonable estimate of firing rate (or Δ*F/F*_0_) as a function of animals’ location. On the other hand, animals would typically not perform more than ∼four trials per test session before becoming satiated. Consequently, the overall estimate of neurons’ tuning curves for different trial conditions is strongly undersampled and contains many zero elements, therefore heavily biasing the measure of spatial information.

In response, we opted for the KSG estimator (Kraskov et al., 2004) to compute the mutual information between location and neuronal responses. The KSG estimator does not rely on binning, and investigations show that it is robust against non-uniformity and undersampled data (Kraskov et al., 2004; Gao et al., 2018). There are two variants to the algorithm and, here, we employ the first variant as described in the original article. Briefly, define the animal’s position and a neuron’s firing rate over time by the vectors ***x*** = [*x*_1_*, x*_2_*,…,x_N_*] and ***λ*** = [*λ*_1_*, λ*_2_*,…,λ_N_*], respectively. For each data point (*x_i_, λ_i_*), we find its *k*-th nearest neighbour, indexed at *n*, and measure the longest distance from the point to this neighbour along the separate axes of ***x*** and ***λ***. That is,

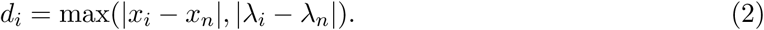

Then, the estimate for the mutual information between ***x*** and ***λ*** (i.e., the spatial information) is

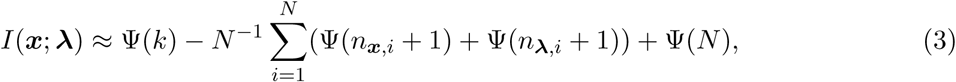

where Ψ(·) is the digamma function, and the counting function

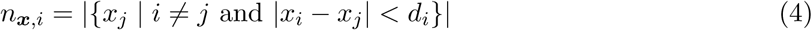

counts the number of data points along the single axis of ***x*** whose distance from *x_i_* is below *d_i_*. The complete algorithm proceeds as follows:

Algorithm 1 KSG mutual information estimator (first algorithm)

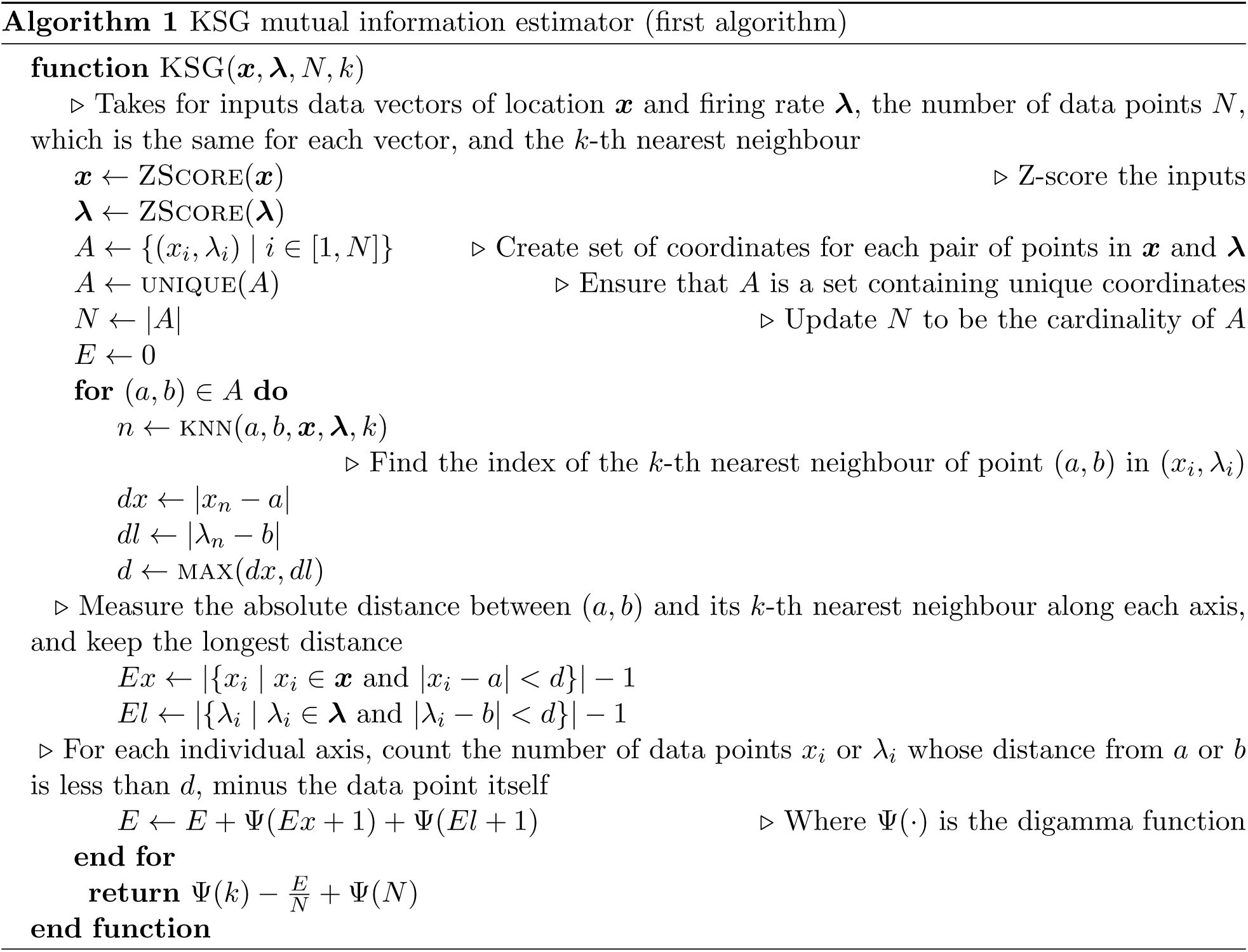

We implemented this algorithm in a custom C library (https://github.com/LelouchLamperougeVI/libKSG) and used it to compute the spatial information for each neuron during the movement epochs. By normalizing the estimates with log 2, we report spatial information in units of bits.

### 6.10 Bayesian decoding

The procedures for Bayesian decoding of animal position from neuronal activity are as previously described (Inayat et al., 2023; Esteves et al., 2023; Demchuk et al., 2024). Briefly, it can be shown that the maximum *a posteriori* estimate of animal position *x*, given a population vector of neuronal activities **n** comprising *N* neurons, is

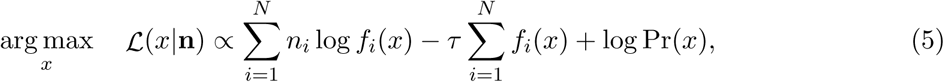

where *n_i_* is the mean activity of neuron *i* in a given time window *τ*, *f_i_*(*x*) is the average activity rate 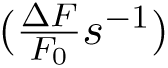 of neuron *i* at position *x*, and Pr(*x*) is the probability of occupancy of position/bin *x*. As such, we work with log-likelihoods as opposed to raw probabilities (cf. Zhang et al. 1998) to avoid the accumulation of floating-point errors from the arithmetic multiplication of many small numbers. In this context, the “training data” is referred to as *f_i_*(*x*) and Pr(*x*), while the “test data” is **n**. We used a *τ* of 500 ms for all decoding.

## 7 Author contributions

ZR: Conceptualization, Data curation, Software, Formal analysis, Validation, Investigation, Visualization, Methodology, Writing - original draft, Writing - review and editing; HC: Software, Formal analysis, Validation, Investigation, Visualization, Writing - review and editing; MHM: Conceptualization, Resources, Supervision, Funding acquisition, Methodology, Writing review and editing; IQW and RJS: Conceptualization, Writing - review and editing

## 9 Supplementary

The supplementary analyses provide histological verification of the imaging site and behavioural controls relevant to interpretation of the CA1 spatial-coding results. Histology confirmed GCaMP8f expression in dorsal CA1 beneath the GRIN lens (Figure S1a). Because neural analyses treated the alley as a linearized environment, positional variance was compared along the longitudinal and lateral axes. Longitudinal variance was approximately 92-fold greater than lateral variance, supporting a one-dimensional representation of position (Figure S1b). We additionally examined whether differences in trajectory geometry could contribute to condition-dependent CA1 coding. Trajectories were characterized using RMS (root-mean-square) lateral deviation, which measures lateral displacement from the direct path, and tortuosity, which measures path directness. These measures were compared among outgoing, EAT, and CARRY runs separately for the short and long alleys (Figure S1c-e). Pairwise comparisons used Wilcoxon signed-rank tests with Holm correction across the 12 comparisons, matched-pairs rank-biserial correlations and median paired differences are reported in (Table S1). No comparison remained significant after correction, suggesting that gross differences in trajectory geometry were unlikely to account for the condition-dependent neural effects.

---

**Figure S1:**
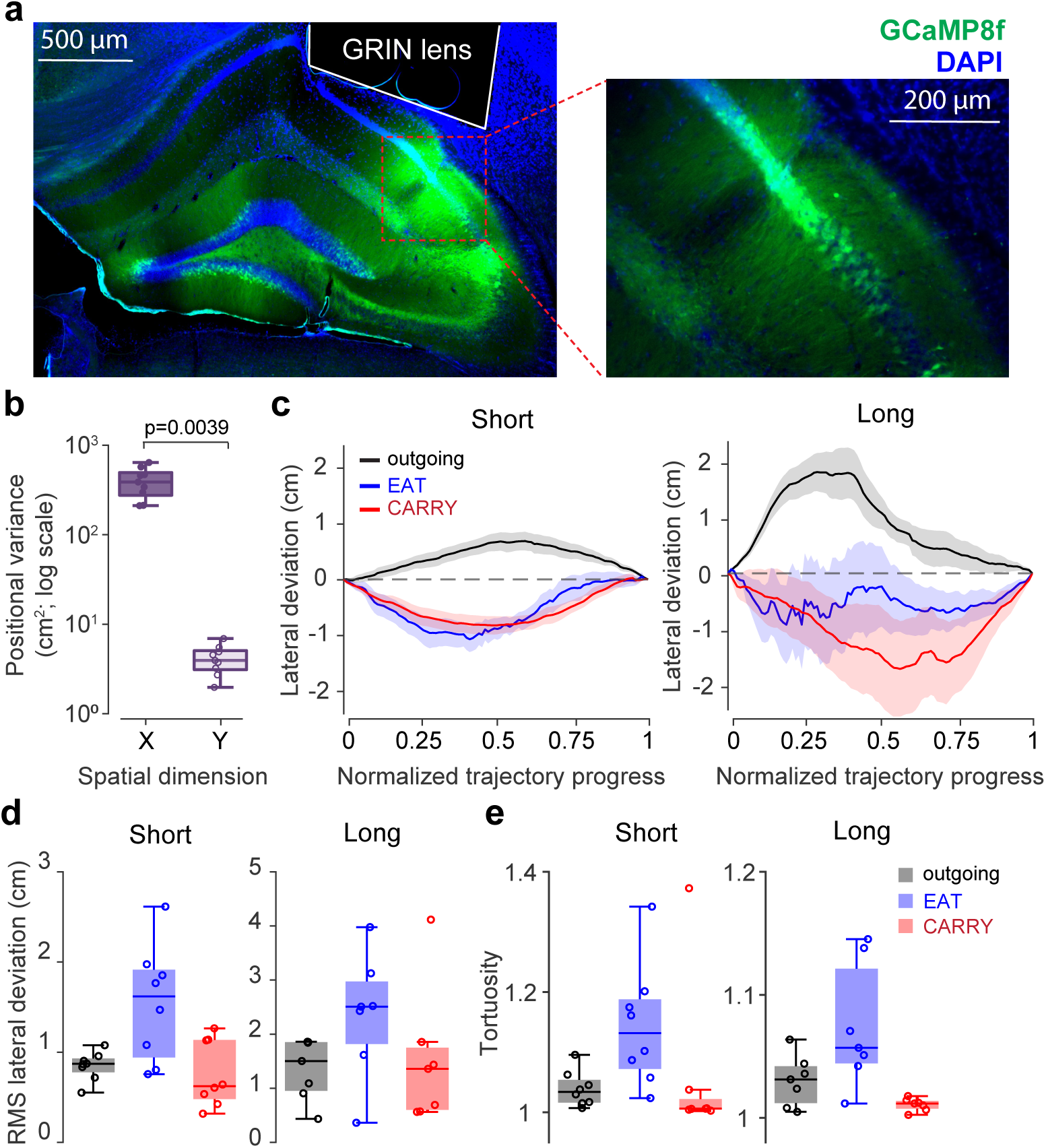
Histological verification and characterization of behavioural trajectories. **a** Representative histological image of dorsal CA1 showing GCaMP expression and a magnified view of the CA1 region beneath the GRIN lens. **b** Positional variance along the longitudinal (along-track) and lateral dimensions of the apparatus. Individual points represent animals, and box plots show the distribution across animals. The y-axis is displayed on a logarithmic scale. **c** Raw trajectories of all trials showing the paths taken by animals between the home and food locations across behavioural conditions. **d-e** RMS lateral deviation and tortuosity of Outward, Eat-inward, and Carry-inward trajectories in the Short and Long conditions. Individual points represent animals, and box plots show the distribution across animals. Pairwise comparisons were performed using paired Wilcoxon signed-rank tests. Because both trajectory measures and both distance conditions were considered complementary components of a single trajectory-control analysis, Holm correction was applied across the complete family of 12 pairwise comparisons. None of the comparisons remained significant after adjustment. However, several comparisons showed large matched-pairs rank-biserial effect sizes, indicating consistent directional differences despite the absence of corrected statistical significance. Raw and Holm-adjusted p-values, rank-biserial effect sizes, median paired differences, and sample sizes for all comparisons are reported in Table S1.

**Table S1:** Pairwise comparisons of trajectory measures across behavioural conditions. Pairwise comparisons were performed using Wilcoxon signed-rank tests. Holm adjustment was applied across the complete family of 12 comparisons. Effect sizes are reported as matched-pairs rank-biserial correlations *r*_rb_). Median differences were calculated as the first condition minus the second condition and are expressed in cm for RMS lateral deviation and are dimensionless for tortuosity. No comparison remained significant after Holm adjustment.

| Distance | Comparison | $p_{raw}$ | $p_{Holm}$ | $r_{rb}$ | Median difference | $n$ |
| --- | --- | --- | --- | --- | --- | --- |
| **RMS lateral deviation** |  |  |  |  |  |  |
| Short | Outward vs. Eat inward | 0.0156 | 0.1719 | -0.944 | -0.644 | 8 |
| Short | Outward vs. Carry inward | 0.4609 | 0.9219 | 0.333 | 0.266 | 8 |
| Short | Eat inward vs. Carry inward | 0.0156 | 0.1719 | 0.944 | 0.608 | 8 |
| Long | Outward vs. Eat inward | 0.0781 | 0.5469 | -0.786 | -1.005 | 7 |
| Long | Outward vs. Carry inward | 0.9375 | 0.9375 | 0.071 | -0.257 | 7 |
| Long | Eat inward vs. Carry inward | 0.0781 | 0.5469 | 0.786 | 1.045 | 7 |
| **Tortuosity** |  |  |  |  |  |  |
| Short | Outward vs. Eat inward | 0.0078 | 0.0938 | -1.000 | -0.081 | 8 |
| Short | Outward vs. Carry inward | 0.1953 | 0.5938 | 0.556 | 0.025 | 8 |
| Short | Eat inward vs. Carry inward | 0.1484 | 0.5938 | 0.611 | 0.088 | 8 |
| Long | Outward vs. Eat inward | 0.0313 | 0.2813 | -0.929 | -0.018 | 7 |
| Long | Outward vs. Carry inward | 0.0781 | 0.5469 | 0.786 | 0.014 | 7 |
| Long | Eat inward vs. Carry inward | 0.0313 | 0.2813 | 0.929 | 0.049 | 7 |

